# Genomic comparisons shed light on the adaptive basis of brain size plasticity and chromosomal instability in the Eurasian common shrew

**DOI:** 10.1101/2025.04.11.648246

**Authors:** William R. Thomas, Tanya M. Lama, Diana Moreno Santillán, Marta Farré, Cecilia Baldoni, Linelle Abueg, Jennifer Balacco, Olivier Fedrigo, Giulio Formenti, Nivesh Jain, Jacquelyn Mountcastle, Tatiana Tilley, Alan Tracey, David Ray, Dina K. N. Dechmann, Dominik von Elverfeldt, John Nieland, Angelique P. Corthals, Erich Jarvis, Liliana M. Dávalos

## Abstract

*Sorex araneus*, the Eurasian common shrew, has seasonal brain size plasticity (Dehnel’s phenomenon) and abundant intraspecific chromosomal rearrangements, but genomic contributions to these traits remain unknown. We couple a chromosome-scale genome assembly with seasonal brain transcriptomes to discover relationships between molecular changes and both traits. Positively selected genes enriched the Fanconi anemia DNA repair pathway, which prevents the accumulation of chromosomal aberrations, and is likely involved in chromosomal rearrangements (*FANCI, FAAP100*). Genes involved in neurogenesis show either signatures of positive selection (*PCDHA6*), seasonal differential expression in the cortex and hippocampus (Notch signaling), or both (*SOX9*), suggesting a role for cellular proliferation in seasonal brain shrinkage and regrowth. Both positive selection and evolutionary upregulation in the shrew hypothalamus of *VEGFA* and *SPHK2* indicate adaptations in hypothalamic metabolic homeostasis have evolved together with Dehnel’s phenomenon. These findings reveal genomic changes central to the evolution of both chromosomal instability and cyclical patterns in brain gene expression that characterizes mammalian brain size plasticity.

**Teaser:** Genomic and expression variations are key to chromosomal instability and seasonal brain plasticity in the common shrew.

## Introduction

*Sorex araneus*, the Eurasian common shrew, exhibits two rare and remarkable phenotypes: extensive intraspecific chromosomal rearrangements (*1*) and Dehnel’s phenomenon, or seasonal size plasticity (*2*–*5*). While chromosomal rearrangements are common across species boundaries, populations of *Sor. araneus* have extensive karyotypic diversity within species, forming distinct chromosomal races (*1*). This shrew also undergoes dramatic, reversible changes in size, including the brain, in response to seasonal environment pressures (*2*–*5*); one of the few mammals known to deploy this wintering strategy (*6*–*8*). *Sor. araneus* reach an initial size maximum as juveniles in their first summer, followed by shrinking of most organs in the autumn, especially the brain. Size reaches a minimum in winter, with rapid regrowth to a second size maximum the following spring (*2, 3, 5*), simultaneous with pubescence for the single breeding season of their disproportionately short lifespan (*9*). Research on the evolutionary basis of these traits is, however, very recent and has been limited to gene expression analyses (*10, 11*).

Previous cellular and molecular investigations of Dehnel’s phenomenon in *Sor. araneus* have discovered that seasonal brain size change occurs without a reduction in neuronal number (*12*), suggesting the evolution of neuroprotective mechanisms. Seasonal gene expression analyses of the *Sor. araneus* hypothalamus showed enrichment of upregulated genes involved in apoptosis regulation and cancer pathways during autumn brain shrinkage, highlighting a balance between cell proliferation and death (*10*). While the anti-apoptotic gene *BCL2L1* was upregulated during shrinkage, *SPHK2*, which promotes apoptosis (*13, 14*) and increases activity in the brains of patients with Alzheimer’s disease (*15*), was evolutionarily upregulated compared to other mammals. These findings pointed toward a finely tuned system that enables common shrews to reversibly regulate brain shrinkage while avoiding the detrimental effects typically associated with neurodegeneration.

Despite those insights, the evolutionary processes involved in the traits that make *Sor. araneus* unique, extensive chromosomal rearrangements and Dehnel’s phenomenon, remain unknown. While intraspecific karyotypic diversity in mammals is not uncommon (*16, 17*), the more than 75 distinct chromosomal races in *Sor. araneus* indicate unique chromosomal instability (*1*).

Similarly, Dehnel’s phenomenon, though most pronounced in the common shrew, is exceptionally rare among mammals, found only in some red-toothed shrews (Soricinae) (*5*), the European mole (*Talpa europaea*) (*6*) and two mustelid species (*Mustela erminea, Mustela nivalis*) (*8*), with a similar phenotype in a domesticated ferret (*Mustela putorius furo*) (*18*) and the pygmy shrew (*Suncus etruscus*) (*7*). Dehnel’s phenomenon is rare, as is the genomic instability of *Sor. araneus*. While the former must be associated with positive selection to adapt to the ecophysiological challenges these species face, the latter could result from relaxed negative selection on genome repair mechanisms.

Dehnel’s phenomenon is hypothesized to have evolved as an adaptive tradeoff between size and metabolic demand during winter (*5*). This strategy is critical for the common shrew, which has one of the highest mammalian basal metabolic rates per unit of body mass (*19*). By reducing the size of energy expensive tissues, shrews decrease the energy required for their maintenance and movement (*3, 11, 20*–*26*). Recent evidence on the metabolic and gene expression changes concurrent with seasonal size change in the common shrew has lent support to this hypothesis (*11*). For example, blood metabolomics and liver transcriptomics indicate increased lipid metabolism and gluconeogenesis during shrinkage (*11*). These may be regulated by adaptive gene expression of the hypothalamus blood brain barrier (*10*). The evolution of efficient metabolic regulation systems is therefore expected during seasonal size change.

A new chromosome-level genome assembly for the common shrew provides an opportunity to investigate how selection has shaped both the convergently evolved Dehnel’s like phenotypes as well as the unique chromosomal instability of these shrews. We compared *Sor. araneus* chromosome rearrangements to a closely related species (*Sun. etruscus*) and applied branch-site models to identify positively selected genes (PSGs) by comparing the common shrew genome to 39 mammalian genome assemblies. To further relate molecular evolution to Dehnel’s phenomenon, we tested for parallel evolution with three other species exhibiting Dehnel’s phenomenon (*Suncus etruscus, Mustela erminea, Mustela putorius furo*) and integrated results with seasonal gene expression data from the common shrew cortex and hippocampus. We discovered gene enrichment of *Sor. araneus* specific PSGs in the Fanconi anemia pathway (*FANCI, FAAP100, PALB2*), associated with DNA repair and longevity, as well as in *VEGFA*, a candidate hypothalamic regulator of metabolism in the common shrew. Genes associated with neurogenesis (*PCDHA6, SOX9*) also showed signatures of parallel evolution across species with Dehnel’s phenomenon, complementing the seasonal changes identified in the hippocampus and cortex linked to cell proliferation (*SOX9*) and Notch signaling. Together, these integrated genomic and expression analyses implicate changes in genes involved in metabolism and cell proliferation in the evolution of Dehnel’s phenomenon, while also characterizing evolutionary processes potentially associated with chromosomal rearrangements and longevity.

## Results

### Genome assembly, annotation and synteny

We sequenced, assembled, and annotated a chromosome-scale reference genome assembly for one male Eurasian common shrew, *Sorex araneus*, following the Vertebrate Genomes Project (VGP) assembly pipeline (*27*) using a combination of Illumina short-read-, PacBio long-read-, and Hi-C sequencing. PacBio sequencing data had 35.71X genome coverage with a genome size of ∼2.40Gb. We used Hi-C chromosomal interactions to further manually curated the genome and were able to anchor >99.8% of the genome to 12 chromosomes (2N=24) (Figure 1A) with a scaffold N50 of 374MB (Figure 1B) and an estimated a genome size of ∼2.23Gb consisting of 73% unique or non-repetitive sequence. By comparison, the previous *Sor. araneus* genome assembly from the Broad Institute consisted of 12,845 unplaced scaffolds (Table1, Figure 1B).

**Table 1.** Chromosome quality of new *Sorex araneus* genome assembly compared to Broad Institute assembly.

|  | Broad Institute (sorAra2) | New assembly (mSorAra2.pri) |
| --- | --- | --- |
| **Total Sequence Length** | 2,423,158,183 | 2,405,993,548 |
| **Total Ungapped Length** | 2,192,103,426 | 2,393,049,584 |
| **Number of Scaffolds** | 12,845 | 65 |
| **Scaffold N50** | 22,794,405 | 374,364,821 |
| **Scaffold L50** | 36 | 3 |
| **Number of Contigs** | 201,420 | 1,492 |
| **Contig N50** | 22,623 | 4,024,507 |
| **Contig L50** | 27,135 | 182 |
| **Chromosomes** | 0 | 12 |
| **Source Location** | Piel Island, United Kingdom | Radolfzell, Germany |

**Figure 1.**
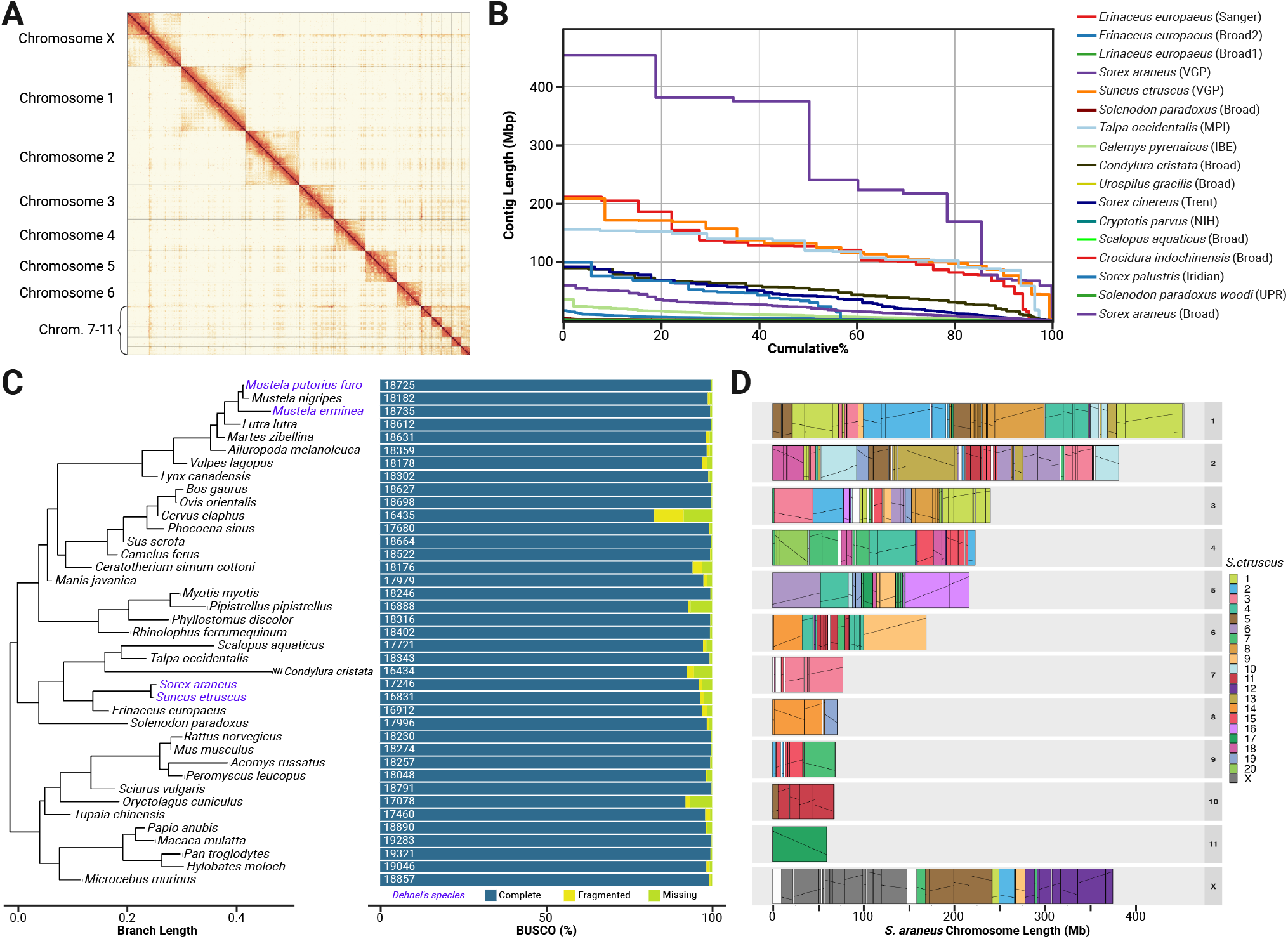
Genome Assembly and Comparisons. **(A)** Using PacBio and Hi-C chromosome interactions and manual curation, the shrew genome assembly was assembled into 12 chromosomes. **(B)** Cumulative percent of each Eulipotyphla genome found on contigs of certain length, showing a highly contiguous shrew genome assembly and longer chromosomes than those of other close relatives. **(C)** Phylogeny of species used in comparative genomic analyses, each with more than 16,000 genes and BUSCO scores greater than 80%. **(D)** Comparisons between *Sor. araneus* and *Sun. etruscus* orthologous blocks. Results highlight orthology between *Sor. araneus* chromosomes 10 and 11 and *Sun. etruscus* chromosomes 11 and 17, as well as fusions of the *Sun. etruscus* X, 5, and 12 and 6, 9, and 16 to form the *Sor. araneus* X chromosome and chromosome 5.

We were not able to recover the Y chromosome. Annotations using TOGA’s machine learning ortholog classifier identified 42,221 transcripts associated with 17,246 genes (BUSCO mammalia_odb10 9,005 complete genes; 97.6%), of which 15,980 were one-to-one orthologs to the human genome (Figure 1C). These numbers are similar to those identified with RNA sequencing, which found 45,362 transcripts associated with 24,205 coding loci, of which 15,752 are annotated genes.

Pairwise alignments between *Sor. araneus* and other species of the order Eulipotyphla with high quality genome assemblies showed conserved syntenic regions between species. Comparing the Eurasian common shrew to its closest relative in the data set, the pygmy shrew *Sun. etruscus*, we found 236 orthologous blocks (Figure 1D). These blocks show that *Sor. araneus* chromosome 10 is orthologous to *Sun. etruscus* chromosome 11, and *Sor. araneus* chromosome 11 is orthologous to *Sun. etruscus* chromosome 17. Pairwise comparisons also show that large portions of *S. araneus* chromosome 7 are syntenic with *Sun. etruscus* 3, and *Sor. araneus* 8 with *Sun. etruscus* 14, while *Sor. araneus* chromosome X appears to be fusions of *Sun. etruscus* X, 5, and 12, with *Sor. araneus* chromosome 5 being fusions of *Sun. etruscus* 6, 9, and 16. Meanwhile, *Sor. araneus* chromosomes 2, 3, and 4 show extensive rearrangements compared to *Sun. etruscus*. Lastly, we inferred the ancestor of Eulipotyphla had 36 chromosomes using DESCHRAMBLER (*28*).

### Positive selection in Sorex araneus

We then analyzed the molecular evolution between the common shrew and other mammal species by testing for positive selection in more than 15,000 single copy orthologs using adaptive branch-site random effects likelihood (aBSREL) models implemented in the HyPhy suite (v2.5.32) (*29*). We inferred 676 genes had been influenced by positive selection (PSGs; p_adj_ < 0.05) in *Sor. araneus* (Figure 2A). Gene set enrichment analysis identified eight Kyoto Encyclopedia of Genes and Genomes (KEGG) pathways enriched by *Sor. araneus* PSGs, consisting of four related to immunity, including complement and coagulation cascades (4.1-fold enrichment, p<0.01), hematopoietic cell lineage (3.1-fold enrichment, p<0.05), primary immunodeficiency (5.1-fold enrichment, p<0.05), and inflammatory bowel disease (3.6-fold enrichment, p<0.05).

**Figure 2.**
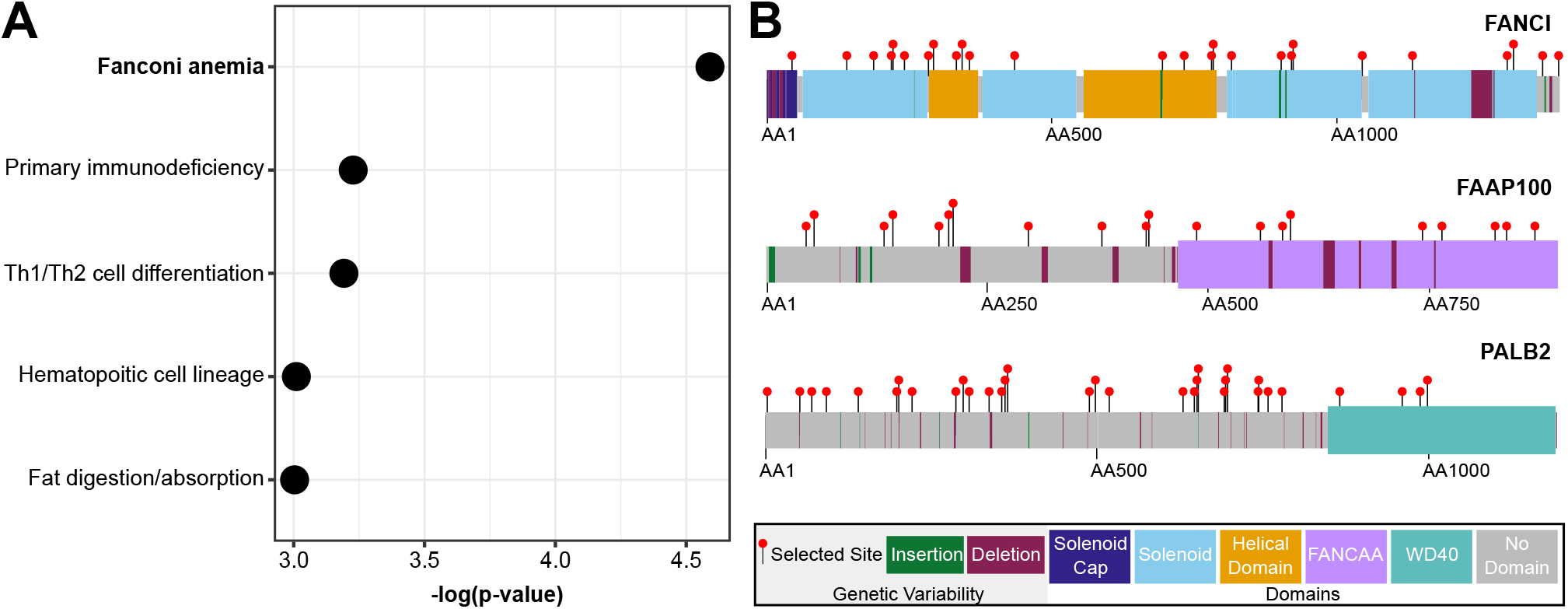
Gene set enrichment of shrew-specific positively selected genes. **(A)** Five pathways were significantly enriched (p<0.05) with positively selected genes only under selection in the *Sor. araneus* lineage. The most enriched pathway was the Fanconi anemia complex, consisting of **(B)** *FANCI, FAAP100*, and *PALB2*, with >19 positively selected sites per gene detected from MEME. Genetic variability (selected amino acids, insertions, deletions) was found across genes, both within and between protein domains.

Additionally, the Fanconi anemia pathway, associated with genomic stability, was enriched (4.3-fold enrichment, p<0.05), including genes *FANCI* (Figure 2), *PALB2, FAAP100*, and *REV3L*. Exploratory analyses on background branches inferred 2997 genes to be evolving under positive selection in at least one background species, of which 288 genes showed signals of pervasive selection throughout the phylogeny (>5 background species). Of the 676 *Sor. araneus* PSGs, 231 genes showed no signal of background selection and were thus specific to this species.

### Parallel evolution of Dehnel’s phenomenon

We tested for selection in more than 10,000 genes annotated in all the selected mammal species with Dehnel-like phenotypes (*M. erminea, M. putorius furo, Sun. etruscus*) and inferred positive selection in over one hundred genes per species, with various degrees of parallel evolution between species. We inferred 225 genes to be under positive selection (p_adj_ < 0.05) in *M. erminea*, 110 genes in *M. putorius furo*, and 904 genes in *Sun. etruscus* (Table 2). Accounting for background selection, eight genes show signatures of parallel evolution with only episodic (<5 species) background selection (*ASPHD1, PCDHA6, TNK2, B9D1, MEGF8, TTLL10, RNF34, LAMTOR4*), significantly more genes than expected by chance (100,000 permutations, p<0.0001), suggesting non-independence with lineage-specific traits, i.e., Dehnel’s phenomenon (Figure 3). A single gene, *PCDHA6*, was parallel between *Sor. araneus, M. erminea*, and *M. putorius furo* with no background selection. MEME analyses were used to detect specific sites evolving under positive selection on *Sor. araneus* PSGs and found several sites to be convergent across species with Dehnel’s-like phenotypes. Across the eight genes inferred to be under parallel evolution, 10 sites were found to have positive selection in multiple species with Dehnel’s-like phenotypes.

**Table 2.** Counts of genes tested and found to be under positive selection by species. For all counts statistical significance at p-val_adj_ < 0.05.

| Species | Total Genes | Count including this tip | Species Specific | Convergent (Shrew+2) | Convergent (<5 Background) | Convergent (<1 Background) |
| --- | --- | --- | --- | --- | --- | --- |
| *Sor. araneus* | 15,812 | 676 | 231 | 23 | 8 | 1 |
| *S. etruscus* | 15,380 | 904 | 397 | 19 | 6 | 0 |
| *M. erminea* | 16,675 | 225 | 54 | 21 | 6 | 1 |
| *M. p. furo* | 16,638 | 110 | 40 | 7 | 4 | 1 |

**Figure 3.**
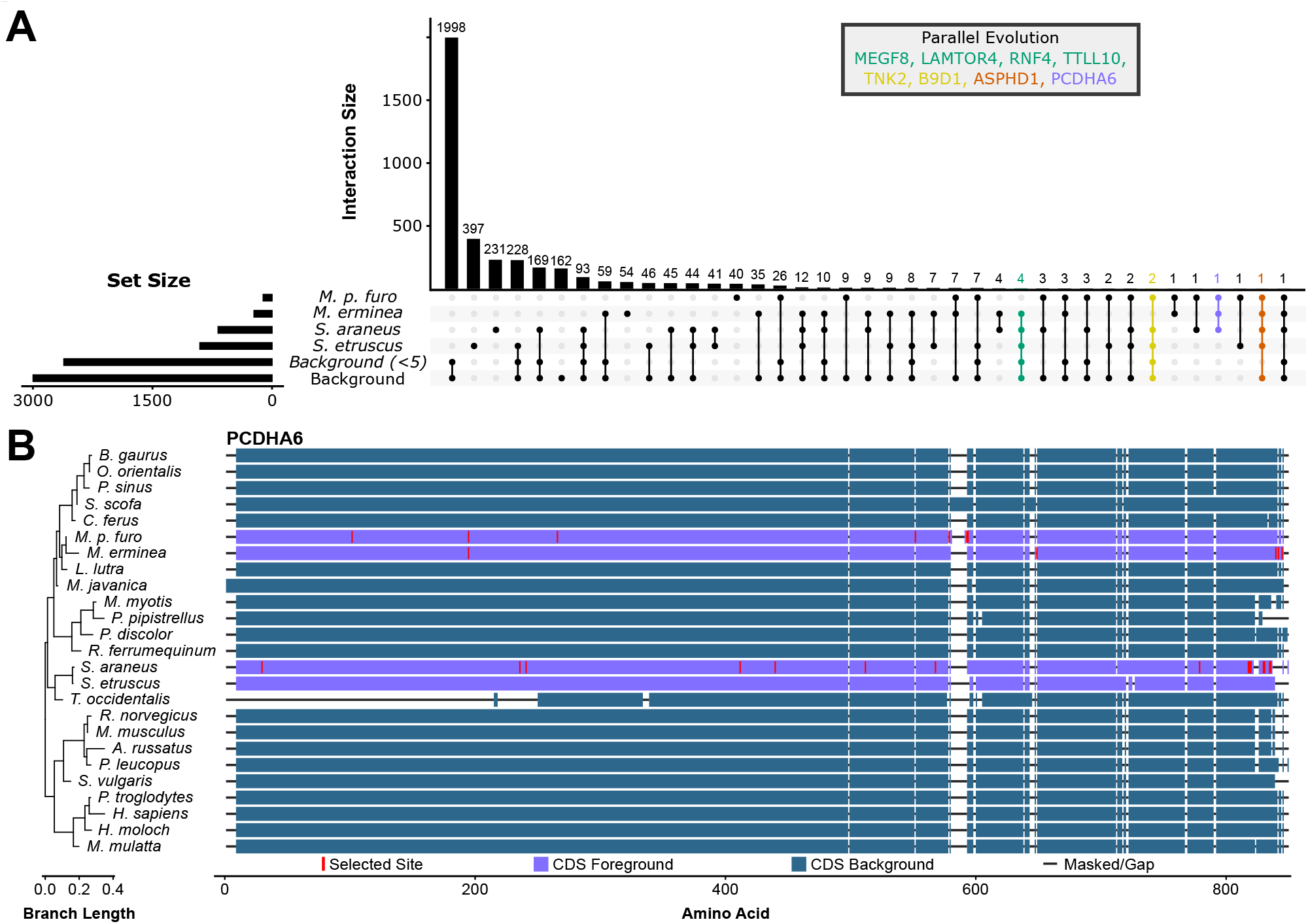
Signatures of parallel evolution in Dehnel’s phenomenon. **(A)** Upset plot of the positively selected genes (PSGs) found in *S. araneus*, species with Dehnel-like phenotypes, and background branches with two degrees of stringency (background selection >0 and background selection >4). Eight genes overlapped across species with Dehnel’s-like traits were found. **(B)** The gene encoding protocadherin alpha 6, *PCDHA6*, was the single gene convergently evolving under positive selection in *S. araneus, M. erminea*, and *M. putorius furo* without background selection.

### Seasonal brain expression

To explore both the regulatory mechanisms of seasonal brain size change and relationships between molecular evolution and function, we tested for differential expression between the shrinking (autumn) and regrowth (spring) phases of Dehnel’s phenomenon in two brain regions, the hippocampus and the cortex, and identified any overlapping PSGs. In the hippocampus, we found 77 differentially expressed genes (DEGs) between autumn and spring, with 52 significantly upregulated in spring and 25 significantly upregulated in autumn (Figure 4). Using a ranked gene set enrichment analysis, we discovered an enrichment of 17 pathways (7 upregulated in autumn, 10 upregulated in spring), which included many pathways associated with metabolism; PPAR signaling (p_adj_<0.01, NES=1.81), fatty acid metabolism (p_adj_<0.01, NES=1.81), and oxidative phosphorylation (p_adj_<0.01, NES=1.81) (Figure 3B). In the cortex, we identified 122 significant DEGs, with 46 upregulated in autumn shrinkage and 76 upregulated in spring regrowth. 10 pathways were enriched with genes varying between these seasons, including five that overlapped in the same direction with those enriched in the hippocampus; ribosome (cortex:p_adj_<0.001, NES=-2.35; hippocampus: p_adj_<0.001, NES=-2.36), protein export (cortex:p_adj_<0.001, NES=-2.31; hippocampus: p_adj_<0.01, NES=-2.05), oxidative phosphorylation (cortex:p_adj_<0.001, NES=-2.02; hippocampus: p_adj_<0.01, NES=-1.78), proteasome (cortex:p_adj_<0.01, NES=-2.05; hippocampus: p_adj_<0.05, NES=-1.67), and Notch signaling (cortex:p_adj_<0.01, NES=1.98; hippocampus: p_adj_<0.05, NES=1.70). Of the DEGs, 6 genes in the cortex and 5 genes in the hippocampus overlapped with PSGs, three of which were found in all three data sets (*SOX9, RIN2, STMN2*) (Figure 5).

**Figure 4.**
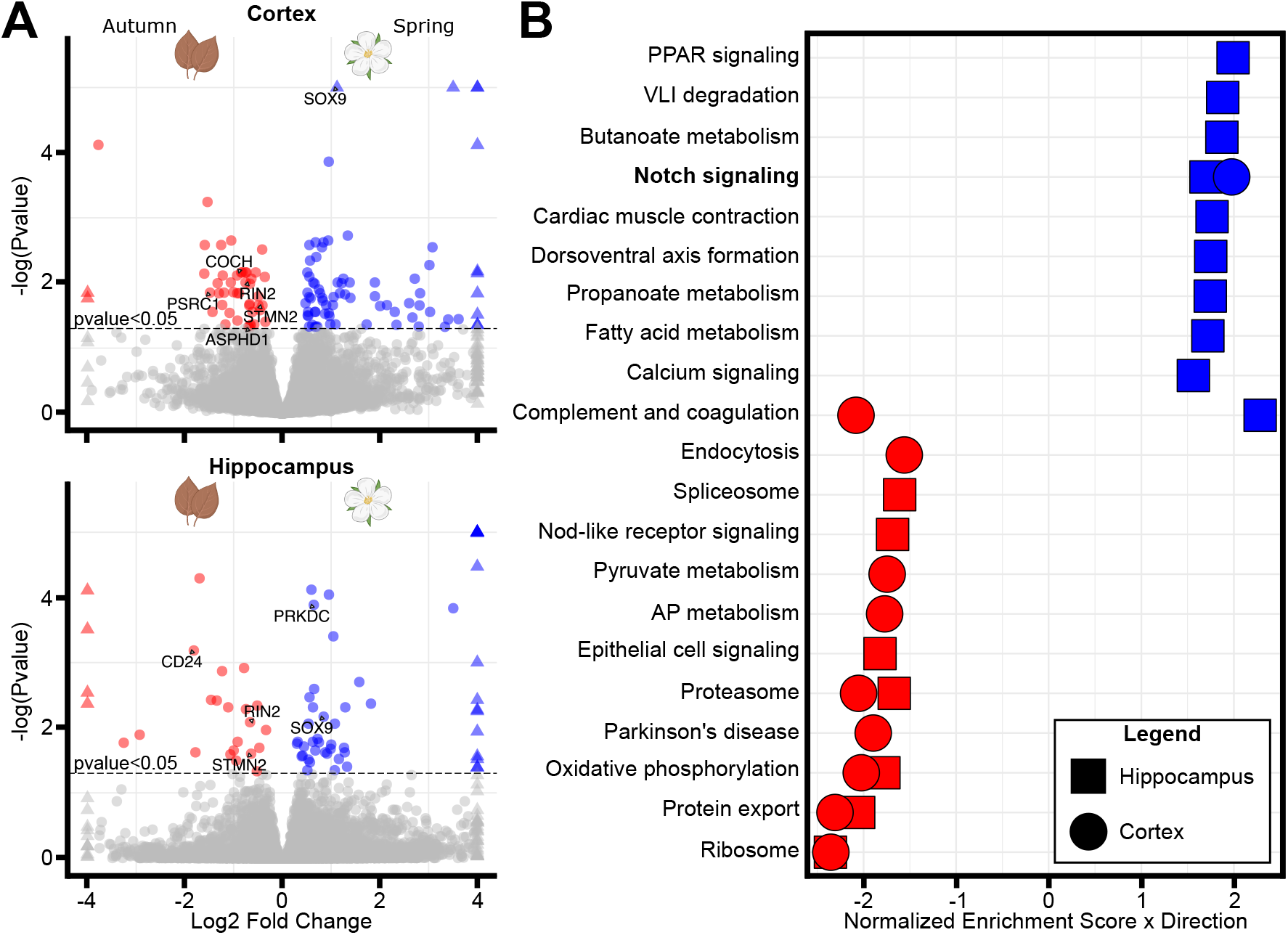
Seasonal gene expression of hippocampus and cortex. **(A)** Volcano plot of differentially expressed genes in the cortex and the hippocampus between autumn (red, cortex: 46 upregulated genes, hippocampus: 25 upregulated genes) and spring (blue; cortex: 76 upregulated genes, hippocampus: 52 upregulated genes). **(B)** Gene set enrichment of differentially expressed genes identified 21 total enriched gene sets (p_adj_<0.05), 5 of which overlap in the same direction between the two brain regions (Notch signaling, proteasome, oxidative phosphorylation, protein export, ribosome).

**Figure 5.**
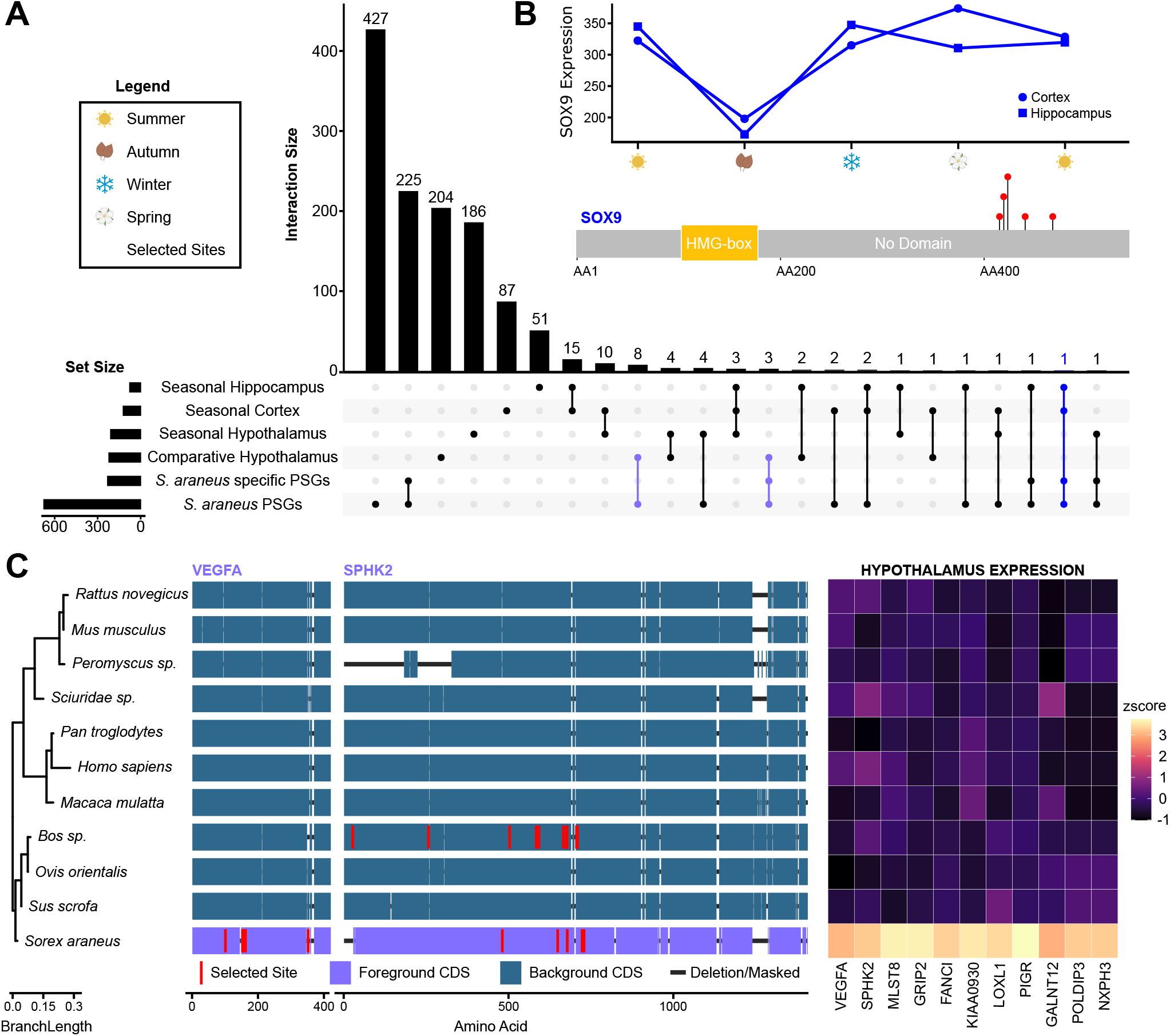
Evolutionary and seasonal expression divergence in the brain associated with positive selection in coding sequence. **(A)** Upset plot of the positively selected genes (PSGs) found in *Sor. araneus* compared to significantly varying genes from transcriptomics analyses. **(B)** *SOX9*, involved in neural stem cell maintenance, is the only gene both downregulated in the hippocampus and cortex during brain shrinkage, and has positively selected changes in its coding sequence outside of a known domain. **(C)** Positive selection was also identified in *VEGFA*, involved in blood brain barrier development and function, and *SPHK2*, which regulates neuronal cell death. Both genes are part of a suite of 11 positively selected genes with upregulated expression in *Sor. araneus* hypothalamus compared to other mammals.

### Hypothalamus expression overlap

There were many instances of overlap between PSGs and prior molecular results. Previous analyses identifying differentially expressed genes in the hypothalamus between seasons of Dehnel’s phenomenon (*10*) only consisted of six overlapping genes with PSGs (*KCNK12, COCH, RSPH3, KNDC1, PARP4, FAN1*). However, comparing PSGs to previous work identifying branch specific change in hypothalamus expression in *Sor. araneus* (Figure 5), we found overlap in genes related to hypothalamus development. Eleven PSGs (*KIAA0930, LOXL1, MLST8, PIGR, VEGFA, GALNT12, POLDIP3, NXPH3, SPHK2, GRIP2, FANCI*) overlapped with those having higher expression in the *Sor. araneus* hypothalamus compared to other mammalian species.

*VEGFA* was found in this overlap which we previously hypothesized was important for *Sor. araneus* hypothalamus fenestration to improve signaling of metabolic demands across the blood brain barrier. Of the eight overlapping genes, three were *Sor. araneus* specific, *LOXL1, GALNT12* and *FANCI*. Overlap between PSGs and functional data may indicate selection in coding sequence to alter gene function and regulation.

## Discussion

### Links between chromosomal evolution, genomic stability, and longevity

Robertsonian chromosomal fusions and rearrangements play a central role in karyotype evolution in the common shrew. This individual *Sor. araneus* had many fewer chromosomes (2N=24) than the pygmy shrew, *Suncus etruscus* (2N=42), indicating many chromosomal fusions in the Eurasian common shrew lineage, or fissions in the pygmy shrew lineage have occurred since their most recent common ancestor (∼20mya) (*30*). Ancestral inference supports fusions in *Sor. araneus*. Pairwise comparisons (Figure 1) between *Sor. araneus* and *Sun. etruscus* show that while some chromosomes remain conserved (*Sor. araneus* C7 ≅ *Sun. etruscus* C3, *Sor. araneus* C10 ≅ *Sun. etruscus* C11, *Sor. araneus* C11 ≅ *Sun. etruscus* C17), many are the result of chromosomal fusions (*Sor. araneus* CX ≅ *Sun. etruscus* CX+ C5+C12, *Sor. araneus* C5 ≅ *Sun. etruscus* C6+C9+C16, *Sor. araneus* C8 ≅ *Sun. etruscus* C14+C19). Evolutionary relationships for many chromosomes are less clear, however. Complicating ancestral inference, *Sor. araneus* has an unusual propensity for chromosomal rearrangements (*31*), with over 75 chromosomal races in a single species (*31*). Based on chromosome number and geography, the closest land races with 11 autosomes are the Cordon and Pelister groups (*31*), but these populations are ∼400 km and 1800 km from the German population, suggesting that this local population may be a distinct chromosomal group.

We found positive selection in genes involved in maintaining genomic stability and their substitutions may explain the evolution of extreme intraspecific chromosomal variation in this species (Figure 2). Chromosome size, arrangement, and number can be disrupted by double-stranded breaks in DNA followed by improper repair (*32, 33*), but evolved mechanisms either prevent such damage or ensure accurate repair. The highly conserved FA gene complex (*34*), comprises 20 genes that identify and repair crosslinked DNA via homologous recombination (*35*).

Several genes associated with this complex were positively selected (six total and four *Sor. araneus* specific PSGs) in *Sor. araneus*. We found evidence of positive selection in *FANCI*, whose encoded protein heterodimerizes with FANCD2 to form an ID complex that localizes to chromatin in response to damage (*35*–*37*). We also identified several positively selected genes related to the FANCI-FANCD2 DNA repair complex: *FAAP100*, which is required for core complex stability with knockouts leading to FA-mediated genomic instability (*38*), and *PALB2*, whose involvement in the PALB2-BRCA2 complex promotes chromatin stability and homologous recombination DNA repair pathways (*39, 40*). Substitutions on this pathway in *Sor. araneus* may impair or alter DNA repair and recombination processes and thus increase the generation of structural variation. As genetic variation provides the substrate for evolutionary change, these rearrangements could also further enhance subsequent local adaptation within this geographically widespread species.

In *Sor. araneus*, substitutions in genes in the Fanconi anemia pathway may be associated with effects of DNA repair on longevity. DNA-damage responses and genomic stability are linked with lifespan evolution, as DNA must be constantly repaired throughout life to reduce the risk of cancer and other age-related diseases (*41*–*44*). Mechanistically, impaired Fanconi anemia DNA repair pathways are linked to premature aging (*45*) and cancers (*46, 47*). Conversely, strong positive selection in DNA repair mechanisms, including *FANCI*, has been found in parrots and other long lived bird species (*48*). In contrast, *Sor. araneus* have a lifespan half that predicted by their size (*9*), reducing the value of maintaining genomic stability. This would relax negative selection on DNA repair pathways and, in turn, co-opt them into alternative functions perhaps related to the propensity for chromosomal rearrangements. But this would also contribute to the short lifespan of *Sor. araneus*, as such modifications would expose older shrews to age-related diseases. Genes within this pathway are excellent candidates for testing for involvement in chromosome evolution of *Sor. araneus* population, as well as longevity, with potential implications for aging research.

### Neuronal proliferation, differentiation, death, and Dehnel’s phenomenon

Distinct molecular changes likely underlie the evolution of Dehnel’s phenomenon, as molecular parallelism associated with these phenotypes was rare. Tests for parallel molecular evolution uncovered eight genes overlapping across species with Dehnel’s-like traits (Figure 3), significantly more than expected by chance (p<0.0001). Consistent with an association with extraordinary brain size plasticity, the gene encoding protocadherin alpha 6, *PCDHA6*, is the only one to show signatures of parallel positive selection across species with Dehnel’s-like phenotypes without background selection. *PCDHA6* is part of the protocadherin alpha cluster, a group of cell adhesion molecules central to brain organization, neuronal circuity, axon maintenance, and synaptic plasticity (*49, 50*). While in mice *PCDHA* knockout leads to miswiring of hippocampal connectivity despite normal neurogenesis (*51*), in cell lines, *PCDHA6* knockout results in altered neurite morphology (*52*). Indicating functional conservation, there was high sequence conservation in species in which this gene was present (15 of 40 assemblies lost or missed the gene). Substitutions in *S. araneus, M. erminea*, and *M. putorius furo* may contribute to maintain, rewire, or regenerate synapses during seasonal changes in brain size and morphology, but *PCDHA6* was not detected in brain expression, weakening the case for the genetic changes found to contribute to brain size plasticity. This dearth of parallel positively selected genes found in Dehnel’s-like phenotypes without background selection makes the common shrew a uniquely important model for researching brain size plasticity.

Comparative tests for convergent evolution did not identify strong candidate genes associated with Dehnel’s phenomenon across species, so we focused on *Sor. araneus* specific PSGs and analyzed gene expression, both seasonally variable and evolutionarily upregulated (the latter previously published) (*10*). While to the best of our knowledge there is no prior study of sequence and expression patterns with the massive changes seen in Dehnel’s phenomenon, the question of how sequence and expression evolve provides crucial context for our results. First, genes expressed in the brain diverge less in their sequences than those expressed in other tissues (*53*).

Second, expression divergence in the brain is also diminished compared to other tissues (*54*). Third, although the rate of nonsynonymous substitutions sometimes correlates with expression divergence (*55*) (e.g., in *Drosophila* (*56*)), this is not always the case especially when analyzed between species (e.g., in birds (*57*)). In short, divergence in either sequence or expression of brain-expressed genes is rare, and the two types of change may or may not be correlated.

Despite the expectation of strongly conserved brain gene expression (*58*), including throughout development (*59, 60*), we identified seasonal expression plasticity in pathways related to cellular proliferation, differentiation, and death. Previously, expression analyses of the *S. araneus* hypothalamus indicated anti-apoptotic pathways, mediated by *BCL2L1* and *NFKBIA*, are upregulated in autumn (*10*). That suggested the common shrew brain actively regulates pathways to avoid cell loss, consistent with the constant or increasing number of neural cells during brain shrinkage (*12*). But we did not observe upregulation of those pathways in the hippocampus or cortex during autumn. Instead, we identified significant changes in Notch signaling in both regions. In *Sor. araneus*, Notch signaling was under expressed during brain shrinkage (Figure 4). Notch signaling is a highly conserved pathway in animals, well characterized for its role in development (*61*) and a master regulator of plasticity in the adult brain of model organisms (*62, 63*).

In the *Sor. araneus* hippocampus and cortex, we found autumn changes in gene expression consistent with enhanced neurogenesis. While the precise effects of the Notch signaling decrease are unknown, decreased Notch can promote neural stem cell proliferation and increased adult neurogenesis (*64*), depleting stem cell populations, and hindering neuronal migration (*65*). In spring, *Sor. araneus* revert to elevated expression of *NOTCH1*, which can decrease the rate of neuron establishment between winter and summer in the hippocampus and cortex (*12*). Notch signaling also regulates *SOX9* expression, which is critical for maintaining neural stem cells (*66, 67*), with *SOX9* knockdown in mice leading to increased neurogenesis (*68*). As with Notch signaling, we observed decreased *SOX9* expression in the cortex and hippocampus of autumn, shrinking *Sor. araneus* (Figures 4 and 5). Consistent with our interpretation of enhanced autumn neurogenesis, winter common shrews increased neuron counts in the hippocampus and cortex compared to summer juveniles (*12*). Despite being highly conserved among background species, as expected for a developmental and brain-expressed gene, *Sor. araneus SOX9* is also under positive selection (Figure 5). Most disease related genetic variation in *SOX9* is associated with developmental disorders, particularly chondrogenesis (cellular bone proliferation) and sex determination (*69*–*71*), however, coding changes could alter the function of this gene in hitherto unsuspected *Sor. araneus* specific neurogenic processes.

### Metabolic homeostasis and Dehnel’s phenomenon

PSGs related to both metabolic regulation and cell proliferation (n=11) also overlapped with genes evolutionarily upregulated in the *Sor. araneus* hypothalamus, supporting the hypothesis that Dehnel’s phenomenon evolved as metabolic adaptation. *VEGFA* and *SPHK2* were under positive selection in *S. araneus* while also showing an approximate four-fold upregulation in the *Sor. araneus* hypothalamus compared to other mammals (Figure 5) (*10*). Upregulation of *VEGFA* can increase blood brain barrier permeability, improving nutrient sensing in the brain, while upregulation of *SPHK2* may be involved with neuronal cell death, as overexpression of *SPHK2* in stressed mice cells promotes apoptosis (*13, 14*). Positively selected substitutions may benefit the common shrew. For example, we propose substitutions in *VEGFA* can alter blood brain barrier vascularization in *S. araneus*, as different isoforms can be pro-or anti-angiogenic (*72, 73*), while substitutions in *SPHK2* may reduce the apoptotic effect of this gene in the brain, allowing evolutionary overexpression of each in the *Sor. araneus* hypothalamus. Future mechanistic experiments can test both functional hypotheses.

### Limitations

We conducted comparative analyses to characterize chromosome evolution and identify positive selection in the protein-coding genes of *Sor. araneus* and other species with Dehnel-like phenotypes, while also examining these results in relation to seasonal and evolutionary changes in brain RNA expression. However, this study has several limitations. To determine if the Radolfzell population is a distinct chromosomal race requires additional cytogenetic verification. While protein coding genes are essential, they are not the sole contributors to phenotypic information within the genome (*74*). A significant portion of the vertebrate genome is non-coding, consisting of Conserved Non-coding Elements (CNEs) and cis-regulatory elements (CREs) that play crucial regulatory roles in gene expression, including in brain development (*75*) and in alternative wintering strategies (*76*). As we explore gene regulation associated with Dehnel’s phenomenon across diverse tissues, future studies should investigate the evolution of common shrew CNEs and CREs, feasible as more chromosome-level genome assemblies of Eulipotyphlans are sequenced. Our findings will benefit from mechanistic validation, as gene function in rodent or primate models may differ from those in the shrew, especially in cases involving coding changes that affect protein structure and function. Perturbation experiments, particularly in candidate genes associated with metabolic regulation and cellular proliferation, will provide future insights into the molecular mechanisms of mammalian brain shrinkage and regrowth.

## Conclusion

*Sorex araneus* has evolved a unique set of traits, including a high rate of chromosomal rearrangements and more seasonal brain size plasticity than any other mammal. Yet, the impacts of both selection on protein-coding genes and gene regulation on these traits were unknown. We therefore generated a chromosome-level genome assembly and conducted comparative genomics and seasonal transcriptomic analyses, with three key findings. First, genes in the Fanconi anemia DNA repair pathway (*FANCI, FAAP100, PALB2*) are under positive selection specific to the *Sor. araneus* lineage. We propose these changes contribute to the Eurasian common shrew’s increased chromosomal rearrangements and reduced lifespan. Second, genes and pathways involved in neurogenesis (*SOX9*, Notch signaling, *PCDHA6*) appear critical to the evolution and regulation of Dehnel’s phenomenon, suggesting processes related to adult neurogenesis may mitigate the negative effects of decreased brain size. Third, genes previously identified as upregulated in the *Sor. araneus* hypothalamus were also under positive selection in *Sor. araneus* (*VEGFA, SPHK2*), highlighting potential synergies between protein-coding and expression adaptations to improve metabolic homeostasis in Dehnel’s phenomenon. Positive selection on sequences is concentrated in pathways involved in DNA repair, metabolic homeostasis, and cellular proliferation, processes we propose are integral to the plasticity of the genome and brain of this species.

## Materials and Methods

### Genome Sequencing and Assembly

A single male European common shrew (*Sorex araneus*) was sampled to generate a highly contiguous reference genome from a population in Radolfzell, Germany (47.9684N, 8.9761 E) under protocols authorized by Regierungspräsidium Freiburg, Baden-Württemberg (35-9185.81/G-19/131). This shrew was euthanized and subsampled into tissues which were immediately flash frozen using liquid nitrogen, and then stored at -80°C until DNA extraction. Nucleic acid extraction, genome sequencing, and assembly were conducted by the Vertebrate Genomes Laboratory (pipeline v2.0). High-molecular-weight (HMW) DNA was extracted from spleen tissue using the Bionano Prep Animal Tissue DNA Isolation Kit according to manufacturer protocols. HMW DNA was used to prepare libraries for three sequencing types from listed kits: 1) SMRTbell Template Prep Kit for PacBio Sequel II HiFi, 2) Bionano Prep Labelling NLRS for Bionano Genomics DLS optical mapping, and 3) Arima-HiC kit for Arima Hi-C v2 chromatin interactions. Following sequencing, initial sequencing metrics were assessed with GenomeScope (*77*). Contigs were generated by assembling and phasing PacBio reads with Hifiasm (v0.15.4) (*78*). Duplications representing heterozygous contigs were identified and reassigned with the purge duplications workflow (v.1.2.5) (*79*). Contigs were then scaffolded using Bionano optical maps with Bionano Solve (v.3.6.1). The assembly was further assembled into chromosomal-level scaffolds with Hi-C chromatin interaction data using SALSA2 (v2.2) (*80*), followed by manual curation for errors with gEVAL (*81*). Assembly metrics were benchmarked throughout the pipeline using Merqury (v1.1) (*82*), Quast (v5.0.2) (*83*), and BUSCO (v3.0.2) (*84*).

### Genome Annotation and Alignment

Gene annotation and orthology inference for *Sor. araneus* and *Mustela nigripes* were predicted with projections from the human genome. First, curated assemblies of these two species were masked for repetitive regions through the creation of a *de novo* repeat library with RepeatModeler (v2.0.2) (*85*) followed by soft masking with RepeatMasker (v4.1.2) (*86*). Masked genomes were then pairwise aligned to the human reference genome (hg38) using lastz (v1.04.15) with the following alignment parameters (BLASTZ_O=400, BLASTZ_E=30, BLASTZ_M=254, chainLinearGap=loose). Sensitivity and specificity of generated pairwise whole-genome alignments were improved using RepeatFiller (v1.0) (*87*) and chainCleaner (*88*). Resultant alignment chains were then used as inputs to predict orthologous gene locations with TOGA (v1.0) (*89*), with annotation completeness for both species assessed with BUSCO v5.2.2 (*84*).

*Sor. araneus* and *M. nigripes* TOGA annotations were then aligned to 38 mammalian TOGA annotations acquired from https://genome.senckenberg.de//download/TOGA/, including the stoat (*Mustela erminea*), the domesticated ferret (*Mustela putorius furo*), and the pygmy shrew (*Suncus etruscus*), all of which also undergo Dehnel-like growth patterns. Aligned species were chosen to reduce overrepresentation of species in orders or superorders (7 Eulipotyphlans, 4 Chiropterans, 6 Artiodactyls, 1 Perissodactyl, 1 Pholidota, 8 Carnivorans, 6 Glires, 6 Primates, 1 Scandentia), with genome annotations by TOGA requiring >16,000 annotated genes with BUSCOs >80% (Supplemental Data). The longest transcript for each gene was selected and aligned using the MACSE exon-by-exon alignment function of TOGA to produce multiple codon alignments and cleaned with TAPER (v1.0.0) default settings (*90*). Species were removed from each codon alignment prior to selection analyses when multiple copies of the gene were present or the alignment contained a frameshift, with codon alignments including at least 60% of the species (>24) retained for selection analyses.

### Selection Models

To infer positive selection for lineages within the phylogeny, we used the adaptive branch-site random effects likelihood (aBSREL v2.2) model (*91*) implemented in HYPHY (Hypothesis Testing using Phylogenies) (v2.5.32) (*29*). First, tree pruning was conducted using ETE Toolkit python wrapper (v3) (*92*), where unrooted trees for each codon alignment were pruned from a larger Bayesian molecular-clock mammalian phylogeny (*93*). Multiple aBSREL tests were run. Branch-site models were used to infer positive selection in genes for *a priori* foreground lineage, including *S. araneus*, and those with Dehnel-like phenotypes (*M. erminea, M. p. furo, S. etruscus*). Then, we conducted an exploratory analysis where all background branches were tested for potential positive selection. This step identified genes evolving under positive selection outside of the focal species to test for lineage specificity. To test if genes significantly rejected the null hypothesis of no positive selection, we conducted a likelihood ratio test (LRT), with p-values quantified using a chi-squared distribution 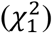 and adjusted for multiple hypothesis tests across genes and species with a Holm-Bonferroni corrections. Running both a priori and exploratory analyses allows identifying genes under positive selection in *Sor. araneus* (regardless of phylogeny-wide selection), genes under positive selection strictly in *Sor. araneus* (no other branches with positive selection), and genes evolving in parallel related to Dehnel’s phenomenon (under positive selection in *S. araneus*, 2 other foreground lineages, <5 other background lineages). To detect potential functional parallelism underlying seasonal size plasticity, we ran a gene set enrichment analysis on our entire candidate gene set using the DAVID functional annotation tool (*94*) with KEGG Gene Ontologies (GO) and Functional Annotations. Finally, we used Mixed Effects Model of Evolution (MEME) (v2.1.2) (*95*) analyses to detect which codons show signals of positive selection per gene using lineages with inferred positive selection. Here, we used a higher threshold for significance (p<0.1), with a stricter Empirical Bayes Factor threshold (EBF>100).

### Chromosomal Evolution

Structural variant analysis of *Sor. araneus* and ancestral karyotype reconstruction of Eulipotyphla were conducted with DESCHRAMBLER (v1) (*28*). Using the new chromosome-scale *Sor. araneus* genome as a reference, we produced pairwise genome alignment chains and nets for five Eulipotyphla species (*Condylura cristata, Galemys pyrenaicus, Suncus etruscus, Talpa occidentalis, Erinaceus europaeus*) and two Chiropterans (*Rhinolophus ferrumequinum* and *Phyllostomus discolor*; outgroups) with high quality assemblies (>5 million scaffold N50, BUSCO > 95%). Chains and nets were made with lastz as described above. Conserved segments, orthology blocks, and predicted ancestral chromosome structure were identified by DESCHRAMBLER with a 1Mb resolution and visualized using syntenyPlotteR (*96*).

### Transcriptomics

Transcriptomic data for the cortex and hippocampus was used to explore the regulatory mechanisms of brain shrinkage and regrowth through Dehnel’s phenomenon and identify further overlap with PSGs. Shrews were sampled, brain regions dissected, and RNA extracted as described in previous studies (*10, 11*). Briefly, common shrews were collected (n=24) across five seasons (n=4-5/season) of Dehnel’s phenomenon (summer juveniles, autumn, winter, spring, summer adults) from a single German population (47.9684N, 8.9761 E) using a protocol that minimized trap-related stress and food insufficiency. Shrews were euthanized with vascular perfusion of PAXgene Tissue Fixative, with brain regions dissected in cold fixative. The cortex and hippocampus were incubated in PAXgene Tissue Stabilizer for 2-24 hours and snap frozen in liquid nitrogen (−180°C). RNA was extracted from each region using a modified Qiagen Micro RNAeasy protocol designed for small amounts of mammalian sensory tissue to reduce heat-associated RNA degradation. Quality control (nanodrop and RNA ScreenTape), library preparation (poly-A selection), and sequencing (approx. 15-25 million reads/sample, 150bp PE) was conducted by Azenta Life Sciences.

For each sample, raw reads were trimmed for adapter sequences and pruned for low quality sequences using fastp (*97*), and then aligned to the novel NCBI *Sor. araneus* transcriptome (GCF_027595985.1_mSorAra2.pri_rna.fna) using Kallisto (v0.46.2) (*98*). Counts were normalized using the DESeq2 (v1.36) (*99*) median of ratios which accounts for library size and content. We then tested for differential expression in both regions between autumn and spring individuals. These seasons were chosen as they resemble the shrinking and regrowth phases of Dehnel’s phenomenon, and were previously analyzed in the hypothalamus (*100*). Genes were tested for differential expression (p_adj_<0.05) using a Wald test in DESeq2 (*99*), with sex as an additional covariate (∼sex + season), followed by Benjamini-Hochberg multiple test correction (*101*). We then ran a KEGG GO enrichment using fsgea (v1.22) (*102*) to describe potential molecular functions or pathways associated with identified differential expression. Lastly, we identified genes that overlapped with genes inferred to be evolving under positive selection.

## Supporting information

Supplementary_Data_AnnotationsInfo

Supplementary_DESeq2

Supplementary_PSG_SorexSpecific

Supplementary_PSG_SorexTotal

Supplementary_Transcriptomics_SampleData

## Acknowledgments

Bioinformatic analyses were conducted on UMass Amherst’s Unity HPC cluster. Additionally, we thank Marion Muturi for support with fieldwork.

## Author contributions

Conceptualization: WRT, LMD

Methodology: WRT, GF, OF, MF, DDMS, TML Software: MF, TML, MF

Validation: WRT Formal analysis: WRT Investigation: WRT Resources: TML

Data Curation: WRT, LA, JB, NJ, JM, TT, AA, CB

Writing – Original Draft: WRT

Writing – Review and Editing: LMD, DD, TML, DDMS, DE, JN, DR, GF, EJ Visualization: WRT, GF

Supervision: LMD, GF, EJ Project administration: LMD

Funding acquisition: LMD, APC, DD

## Data and materials availability

All data are available in the main text or the supplementary materials. Cleaned whole-genome alignments, cleaned gene alignments, and selection outputs openly available on Dryad https://doi.org/10.5061/dryad.8gtht770x. Supplementary tables, results, and code deposited and found on Github https://github.com/wrthomas315/Sorex_Genome2. Raw sequencing data located in the NCBI Sequencing Read Archive (BioProject PRJNA941271). Biorender was used to generate Figure 1 and ChatGPT was used to improve manuscript grammar.

